# Insight into the ecology of vaginal bacteria through integrative analyses of metagenomic and metatranscriptomic data

**DOI:** 10.1101/2021.06.17.448822

**Authors:** M. T. France, L. Fu, L. Rutt, H. Yang, M. Humphrys, S. Narina, P. Gajer, B. Ma, L. J. Forney, J. Ravel

**Affiliations:** Institute for Genome Sciences and Department of Microbiology and Immunology, University of Maryland School of Medicine, Baltimore, MD USA; Institute for Bioinformatics and Evolutionary Studies and Department of Biology, University of Idaho, Moscow, ID USA

**Keywords:** Vaginal microbiome, microbial ecology, metatranscriptome, metagenome, host-microbe interactions

## Abstract

Vaginal bacterial communities dominated by *Lactobacillus* species are associated with a reduced risk to various adverse health outcomes. However, somewhat unexpectedly many healthy women have microbiota that are not dominated by lactobacilli. To determine the factors that drive vaginal community composition we characterized the genetic composition and transcriptional activities of vaginal microbiota in healthy women. We demonstrated that the abundance of a species is not always indicative of its transcriptional activity and that impending changes in community composition can be predicted from metatranscriptomic data. Functional comparisons highlight differences in the metabolic activities of these communities, notably in their degradation of host-produced mucin but not glycogen. Degradation of mucin by communities not dominated by *Lactobacillus* may play a role in their association with adverse health outcomes. Finally, we show that the transcriptional activities of *L. crispatus, L. iners*, and *G. vaginalis* vary with the taxonomic composition of the communities in which they reside. Notably, *L. iners* and *G. vaginalis* both demonstrated lower expression of their cholesterol-dependent cytolysins when co-resident with *Lactobacillus* spp. and higher expression when co-resident with other facultative and obligate anaerobes. The pathogenic potential of these species may depend on the communities in which they reside and thus could be modulated by interventional strategies. Our results provide insight to the functional ecology of vaginal microbiota and reveal strategies for management of these ecosystems.

## Background

The microbial communities that inhabit the human body are crucial determinants of health (1-3). Of these, the microbiota that inhabit the human vagina are unique (4, 5). They dramatically differ in taxonomic composition from the communities found in the vaginas of other mammals, including non-human primates (6), and often have a skewed structure in which the communities are dominated by a single species of *Lactobacillus* (7-11).

The vaginal microbiota in reproductive age women can be roughly divided into five distinct community state types (CSTs) based on species composition (7, 9). Four of these are characterized by a high relative abundance of different *Lactobacillus* species (CST I-*L. crispatus* dominated, CST II-*L. gasseri* dominated, CST III-*L. iners* dominated, CST V-*L. jensenii* dominated), while the fifth (CST IV) is characterized by a more diverse assemblage of facultative and obligate anaerobes. Studies have concluded that women with communities dominated by *Lactobacillus* spp. are less likely to experience a number of adverse health outcomes, supporting a protective role for these species (12-20). Lactobacilli are thought to provide protection through their production of lactic acid which acidifies the environment, although other factors are likely to contribute as well (21-25). Women whose microbiota are largely depleted of lactobacilli are at greater risk to symptomatic bacterial vaginosis (BV), sexually transmitted diseases, and preterm birth (26-30). Interestingly, the vaginal microbiota has also been shown to sometimes exhibit changes in composition through time, sometimes demonstrating dramatic and rapid changes from being dominated by *Lactobacillus* spp. to CST IV-like communities (31, 32). The factors that drive these patterns are not currently understood.

Most studies of vaginal microbiota have relied on amplicon sequencing of variable regions of 16S rRNA genes to assess community composition, although a growing number of studies have employed shotgun sequencing of the vaginal metagenome (33-38). Only a handful of studies, with few subjects, have examined the vaginal metatranscriptome. For example, a small study by Macklaim, Fernandes (39) focused on the transcriptional activity of *L. iners* in four reproductive age women and found variation in the species’ transcriptional activity. A more recent study examined the metatranscriptome of 14 reproductive age women diagnosed with BV and treated with metronidazole (40). The authors found a set of *Gardnerella vaginalis* CRISPR-associated Cas genes that were upregulated in patients who did not respond to treatment. Finally, a recent study examined the metatranscriptome of over 100 pregnant women in order to identify microbiota driven causes of preterm birth (PTB, (36)). They found a number of genes whose expression was associated with either PTB or term birth, including a number of genes encoding bacterial secretion systems or predicted secreted proteins. A concurrent study from the same group described the vaginal composition and transcriptional activity in pregnant women from a number of racial-ethnic backgrounds (37) and reported a number of pathways associated with nucleotide metabolism whose expression was lower in women of African descent. Although the results of these studies have been illuminating, much remains to be learned from analyzing the vaginal metatranscriptome. Fine scale, integrative analyses of these data have the potential to derive translational *in vivo* mechanistic understanding of the role of the vaginal microbiome in health and disease by characterizing interactions between community members and between the microbiota and host tissues.

Here, we present an analysis of 195 metagenomic and 180 metatranscriptomic datasets from 39 non-pregnant reproductive age women who self-collected vaginal swabs daily over the course of 10 weeks (41). These women self-identified as belonging to a number of different racial backgrounds and were found to have vaginal microbiota classified to different CST. We perform integrative analyses of these data in order gain insight into the functional ecology of the vaginal microbiota.

## Results

### Metagenomic and metatranscriptomic datasets

We characterized the vaginal microbiota of 39 North American reproductive age women (age range: 19-45). Subjects included in the study self-identified as Black or African American (n=24), White or Caucasian (n=10), Hispanic or Latino (n=4), and Asian (n=1). For each subject we chose up to five sampling times that were spaced approximately two weeks apart. The metagenomes (n=195) and metatranscriptomes (n=180) of these samples were sequenced and the resulting reads were mapped to VIRGO, a non-redundant and comprehensive gene catalog of the vaginal microbiome (34). This reference-based approach allowed us to estimate the abundance and expression levels of genes and facilitated the downstream integration of metagenomic and metatranscriptomic data. The mapping results were normalized for gene length and then used to determine the taxonomic composition of both the metagenome and metatranscriptome (Figure 1). The composition of a metagenome represents the relative abundances of taxa, whereas the composition of the metatranscriptome represents the relative transcriptional activities of taxa (Supplemental Table 1).

**Figure 1:**
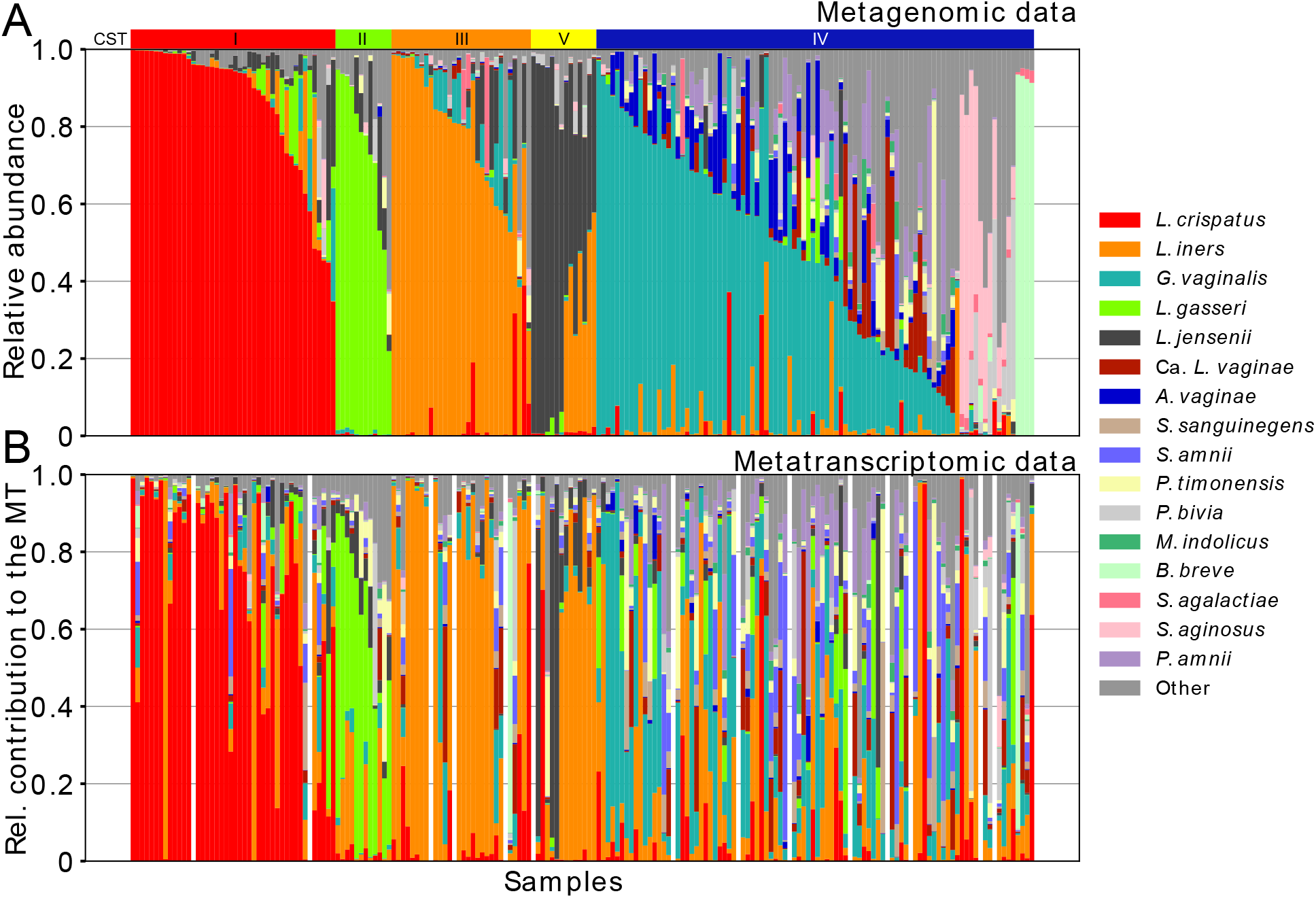
Taxonomic composition of the vaginal microbiota of 39 women at up to five timepoints as assessed by shotgun metagenomics (n=194, panel A). Relative contribution of each species to the metatranscriptome of the corresponding samples (n=180, B). Both estimates were derived VIRGO mapping results and were corrected for gene length. Samples were sorted in the same order in the two datasets, gaps indicate the fourteen samples for which no corresponding metatranscriptomic data were available. Communities were assigned to CSTs based on their taxonomic composition using VALENCIA (9) according to the following scheme: CST I –*L. crispatus* dominated, CST II–*L. gasseri* dominated, CST III–*L. iners* dominated, and CST V–*L. jensenii* dominated) and CST IV. Separate plots displaying the data for each individual subject can be found in Supplemental File 1.

### The relative abundance of a species is not necessarily indicative of its contribution to the metatranscriptome

We first determined whether the relative abundance of a species was consistent with its relative transcriptional activity. The ratio of a species relative activity divided by its relative abundance was referred to as its “relative expression” and represents the extent to which the species is over- or under-represented in the metatranscriptome, as compared to the metagenome. We calculated these relative expression values for the twenty-five most prevalent species study-wide which included the four common vaginal *Lactobacillus* species, *G. vaginalis, Atopobium vaginae, Ca*. Lachnocurva vaginae, and *Prevotella* spp., as well as several other facultative or obligate anaerobic bacteria (Figure 2A). For each species and across samples, a wide range of relative expression values were observed ranging from over- to under-expressive. *G. vaginalis, A. vaginae, Finegoldia magna, Mobiluncus mulieris*, and *Streptococcus anginosus* were more likely to be under-expressive (median relative expression <0, Figure 2A), while the other twenty species were more likely to be over-expressive (median relative expression >0), especially *Sneathia amnii* (median relative expression = 3.3, Figure 2A) and *S. sanguinegens* (median relative expression = 2.2, Figure 2A).

**Figure 2:**
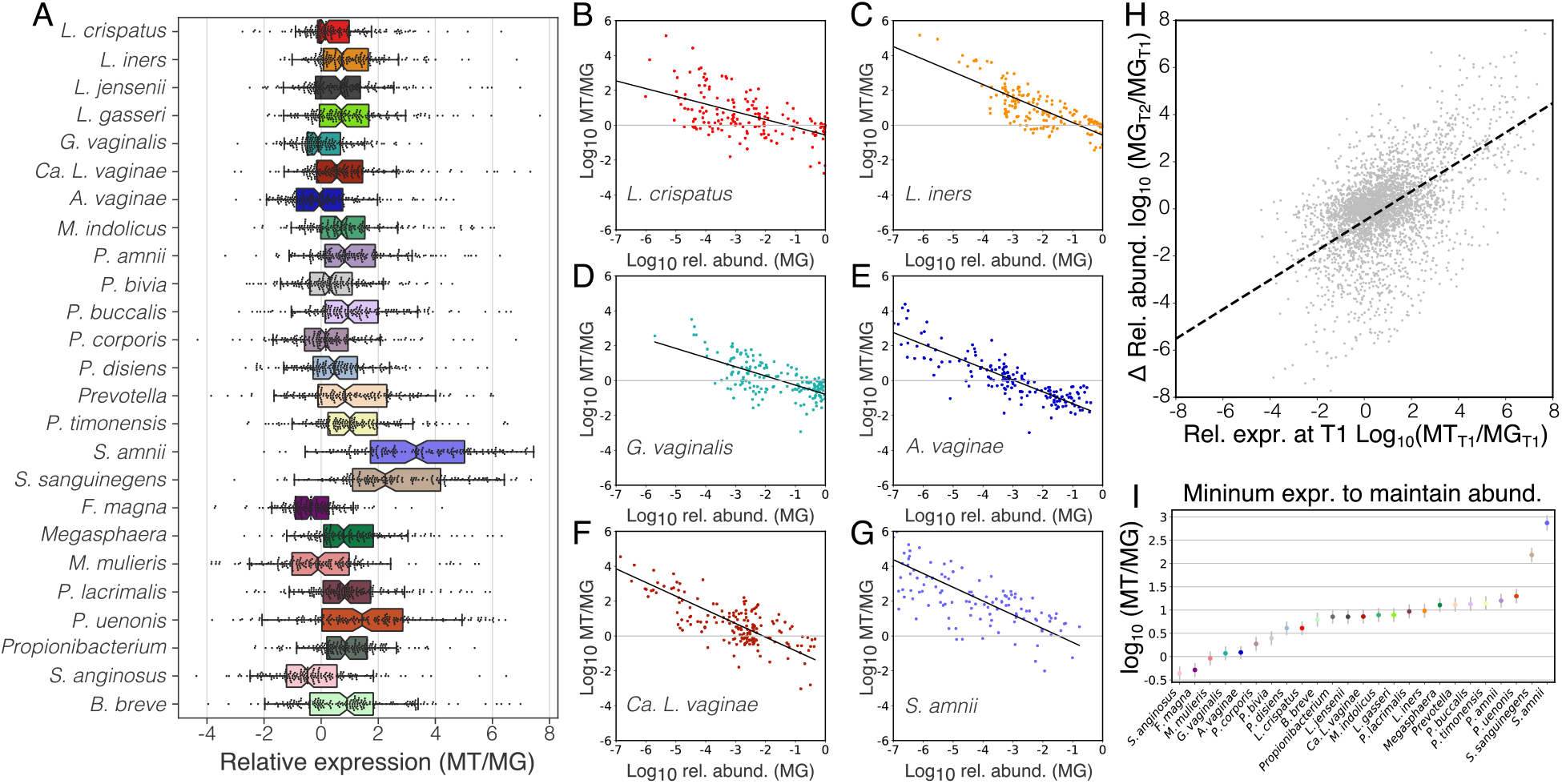
Relative gene expression in the 25 most abundant species in vaginal communities calculated as a species contribution to the metatranscriptome divided by the its relative abundance in the metagenome (A). Relationship between a species’ relative gene expression and its relative abundance in a community (B-G, Supplemental Figure 1). Log_10_ fold change in the relative abundance of a species from one timepoint (T1) to another (T2) as a fuction of its relative gene expression at T1 (H). Minimum relative gene expression requred for a species to maintain its relative abundance from T1 to T2 (I).

We next asked whether a species’ observed relative expression was correlated with its’ relative abundance. Because both datasets are compositional, our evaluation of relative expression was bounded at the highest and lowest relative abundances. If we estimate a species’ relative abundance to be 90%, the maximum value for its relative expression is ∼1.1. Conversely, if a species comprises a small fraction of the community, the minimum value for its’ relative expression is constrained by the depth at which the metatranscriptome was sequenced. We found a strong relationship between the abundance of a species and its’ relative expression (Figure 2B-G, Supplemental Figure 1; linear mixed model; F_1,4261_=7167, p<0.001). When rare, vaginal bacteria often contribute more to the metatranscriptome than their relative abundance would suggest. Whereas species found at higher relative abundances tend to under contribute to the metatranscriptome. The relative abundance at which this switch from over- to under-expressive occurs varied by species (Figure 2B-G; Supplemental Figure 1; linear mixed model; F_24,4261_=198, p<0.001). Species such as *L. crispatus, L. iners*, and *G. vaginalis* (Figure 2B-D), which tend to be found at higher relative abundances, maintain over-expression when common. Conversely, species like *A. vaginae* and *Ca*. L. vaginae (Figure 2E,F), which tend to be found at moderate or low relative abundances, transition to under-expression at lower relative abundances. *S. amnii* (Figure 2G) and *S. sanguinegens* (Supplemental Figure 1) are both routinely estimated to constitute a small proportion of the community, yet they vastly over-contribute to the metatranscriptome.

### A taxon’s contribution to the metatranscriptome may predict future changes in its abundance

The relative transcriptional activity of a species or strain may have some bearing on its future abundance in the microbial community. One might expect that species which are under expressive may experience a decrease in relative abundance, while those that are over expressive may increase. Our longitudinal dataset allowed us to test this hypothesis. For each subject, we have data from up to five timepoints, spread two weeks apart and were able to assess the change in the relative abundance of species between pairs of consecutive timepoints. These values were log_10_ transformed and modeled in response to the species, the log_10_ transformed relative expression of the species and their interaction. As hypothesized, we found a strong relationship between a species’ change in relative abundance and its level of expression at the earlier timepoint (Figure 2H; linear mixed model; F_1,1847_=649, p<0.001). Species that were under-expressive were found to decrease in relative abundance while those that were over-expressive were found to increase. We also find support for species specific intercepts in this relationship (Figure 2I; linear mixed model; F_24,1847_=11.1, p<0.001). These intercepts represent the relative expression necessary for the species to maintain its relative abundance over the two-week interval. We found that most taxa need to be slightly over-expressive to maintain their relative abundance. Notable exceptions were *S. anginosus* and *F. magna* which were found able to maintain their relative abundance while being under-expressive, and *S. amnii* and *S. sanguinegens*, which needed to be highly expressive to maintain their relative abundance.

### Functional differences in the transcriptional activity of communities assigned to different CSTs

By definition CSTs have substantial differences in composition and consequently it is expected that there will also be substantial differences in their functional capacity. To characterize these differences, we performed a differential expression analysis at the level of KEGG orthologs, which represent molecular functions (42). We focused our analysis on comparisons of CST I, CST III and CST IV, as there were few examples of CST II and V in this dataset. Comparisons between these CSTs revealed a large number of differentially expressed KEGG orthologs (Figure 3A-C, full table in Supplemental Table 2). These included expected differences like a 4.6 and 4.8-fold higher expression of D-lactate dehydrogenase (K03778) in CST I compared to CSTs III and IV, which catalyzes the production of D-lactate from pyruvate. The number of differentially expressed KEGG orthologs and the magnitude of those differences was highest between CST I and IV (Figure 3B). The majority of the identified differentially expressed KEGG functions where also differentially abundant in the metagenomic data, signifying an underlying disparity in the functional capacity of these communities. However, there were some KEGG orthologs which were equally abundant but differentially expressed (denoted by yellow diamonds in Figure 3A-C). For example, the *ptsAB* phosphate transport system (K02036 & K02038), which had an equal abundance in CSTs I, III, and IV, exhibited an ∼4-fold higher expression in CST I versus CST III and IV. Another example is alpha-L-rhamnosidase (K05989), which had equal abundance in CSTs I and IV, but 3-fold higher expression in CST IV.

**Figure 3:**
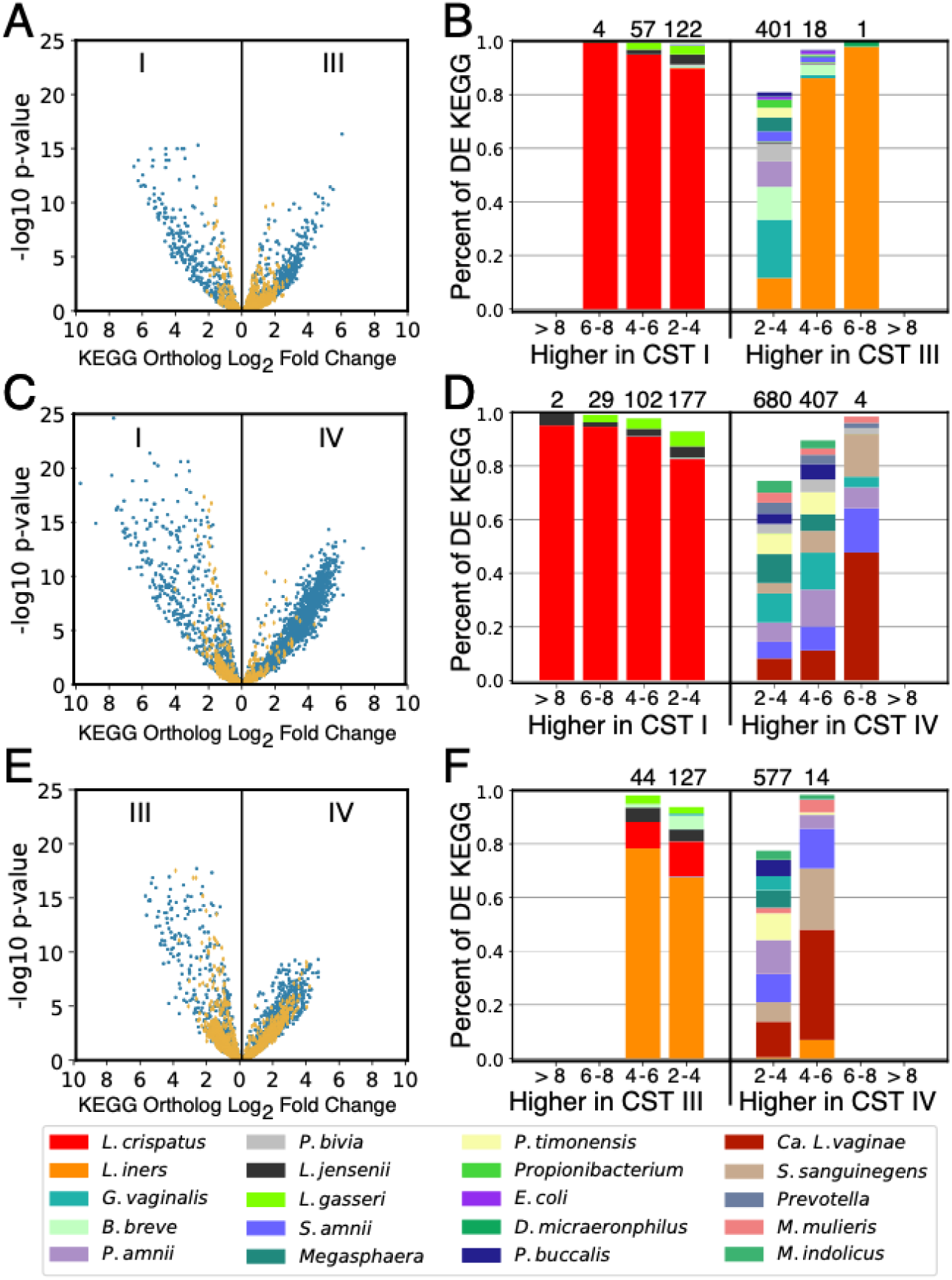
Volcano plots displaying log_2_ fold change differences in the expression of KEGG orthologs between communities assigned to different community state types (A-C). Points in blue represent KEGG orthologs which were identified as differentially expressed and differentially abundant. Points in yellow represent KEGG orthologs that were found to be differentially expressed but not differentially abundant. In panels D-F the taxa responsible for the expression of these KEGG orthologs are displayed. For each fold change category, the percent of each taxa contribution is displayed, averaged across all of the KEGG orthologs in that category. Numbers above the bars indicate the number of differentially expressed KEGG orthologs in the category.

We next characterized the taxa contributing to each of the differentially expressed KEGG orthologs in order to identify the drivers of functional variation in the activity of CSTs (Figure 3D-F. This analysis was enabled by the pairing of taxonomic and functional annotations attributed to genes in the VIRGO database (34). For each KEGG ortholog we calculated the average contribution of each taxon to its expression in each CST. As expected, we found that KEGG functions that are enriched in CST I, compared to either CST III or CST IV, originate mostly from *L. crispatus* (Figure 3D,E). This is not surprising given that communities assigned to CST I characteristically contain a large proportion of this species. In contrast, functions that are enriched in CST III compared to CST I do not primarily originate from *L. iners*, the species which defines CST III. Instead, the expression of these functions primarily originated from species like *G. vaginalis, Bifidobacterium breve*, and several *Prevotella* spp. (2 to 4-fold difference, Figure 3D). This likely reflects the tendency of *L. iners* to coexist with these taxa. Only KEGG orthologs with higher fold changes in this comparison can be primarily attributed to *L. iners*. Expression of KEGG orthologs which were found to be enriched in CST IV compared either to CST I or CST III, were attributed a diverse array of species. *G. vaginalis* and *A. vaginae*, two of the taxa characteristic of communities assigned to CST IV, are not responsible for a majority of these differentially expressed KEGG orthologs; *Ca*. L. vaginae, *Sneathia* spp. and *Prevotella* spp. instead express these functions.

Differentially expressed KEGG orthologs were also grouped into KEGG modules. This higher-level of functional classification describes KEGG orthologs which act in concert to perform a function. Hierarchical clustering was used to group the KEGG modules based on their pattern of associations with CSTs I, III, and IV. This distinguished seven groups of KEGG modules (Figure 4). One group included KEGG modules that demonstrated higher expression in CST I than in CST III or IV. This group includedβ-glucoside, mannitol, phosphonate, phosphate, and putrescine transport systems, as well as the biosynthesis of cysteine, lysine, carotene, and coenzyme F_420_. Another group of KEGG modules was associated with higher expression in CSTs I and III and included: Zn/Mn, glutamine, glucitol transporters, as well as the arbutin-like PTS system. There was no group of KEGG modules which demonstrated exclusively higher expression in CST III, likely reflecting *L. iners* limited functional capacity resulting from its reduced genome size (43, 44). Instead, there were two groupings of modules which demonstrated higher expression in CST III and IV. The association with CST III of these modules is mostly driven by organisms other than *L. iners*. Included within these two groups are: glutamate transport, methionine degradation, iron and zinc transport, and histidine biosynthesis. Finally, there is a number of modules associated only with CST IV. This group includes branched chain amino acid transport, maltose/glucose transport, cytolysin transport, chemotaxis, galactosamine transport, LPS transport, and sulfur degradation.

**Figure 4:**
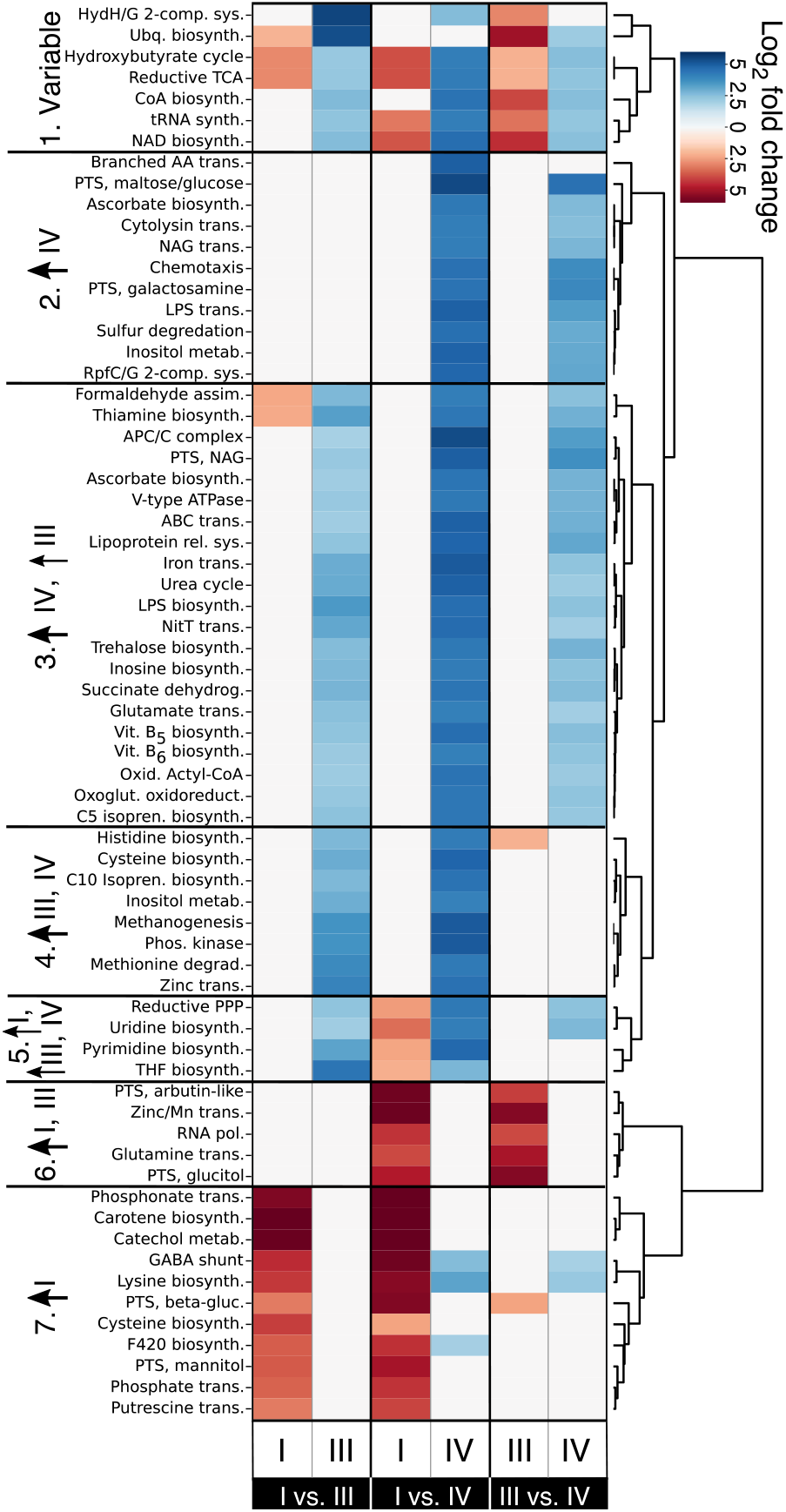
Differentially expressed KEGG orthologs were mapped to KEGG modules, representing metabolic functions. Heatmap displays the average log_2_ fold change values for KEGG modules which demonstrated at least a four-fold difference in one of three comparisons (CST I versus CST III; CST I versus CST IV, or CST III versus CST IV). Two columns are plotted for each of the three comparisons, with each representing KEGG modules whose orthologs demonstrated higher expression in the specified CST. For some KEGG modules which included KEGG orthologs that had higher expression in each of the compared CSTs. Hierarchical clustering was performed on the fold change values using Euclidean distance and Ward linkage to separate the modules into seven categories.

### Expression of glycogen and mucin utilization genes by the vaginal microbiota

Host-produced glycogen and mucin are both believed to be important sources of carbon and energy for bacteria in the vagina (45, 46). Recent work has identified the genes responsible for the metabolism of these products by the microbiota including glycogen debranching enzymes in *Lactobacillus* spp. (47, 48) and sialidases in *G. vaginalis* (49). We examined these and other enzymes involved in the degradation of glycogen and mucin to determine if their expression varied across communities assigned to different CSTs. VIRGO genes responsible for these activities were identified using the Carbohydrate Active enZymyes (CAZy) annotation schema (50). Enzymes placed in the glycosyl-hydrolase (GH) family 13 and glycosyl-transferase (GT) family 35 were identified as likely having glycogen debranching and glycogen phosphorylase activities, respectively. The debranching enzymes act on α-glycoside linkages and can liberate trehalose, glucose, maltose, and maltotriose from glycogen. Expression of these enzymes was similar across the CSTs occurring at around 1,250 transcripts per million (TPM), although it was slightly higher in CST III communities (linear mixed model; Figure 5A). Glycogen debranching enzymes are encoded in the genomes of many bacteria common to the vagina including *Lactobacillus* spp., *G. vaginalis, Ca*. L. vaginae, *Prevotella* spp., and *Sneathia* spp. The expression of glycogen debranching enzymes was consistent with the species composition of CSTs. For example, *L. crispatus* contributes the most to this activity in CST I (Figure 5B). The expression of glycogen phosphorylase enzymes, which act to cleave individual glucose monomers from glycogen, did vary by CST with their expression being highest in CST IV, and lowest in CST I (linear mixed model, Figure 5C). This is likely due to the absence of these genes in the genomes of *Lactobacillus* spp. which are dominant in CST I and III. The taxa mostly responsible for the expression of glycogen phosphorylase enzymes was similar across CSTs (Figure 5D).

**Figure 5:**
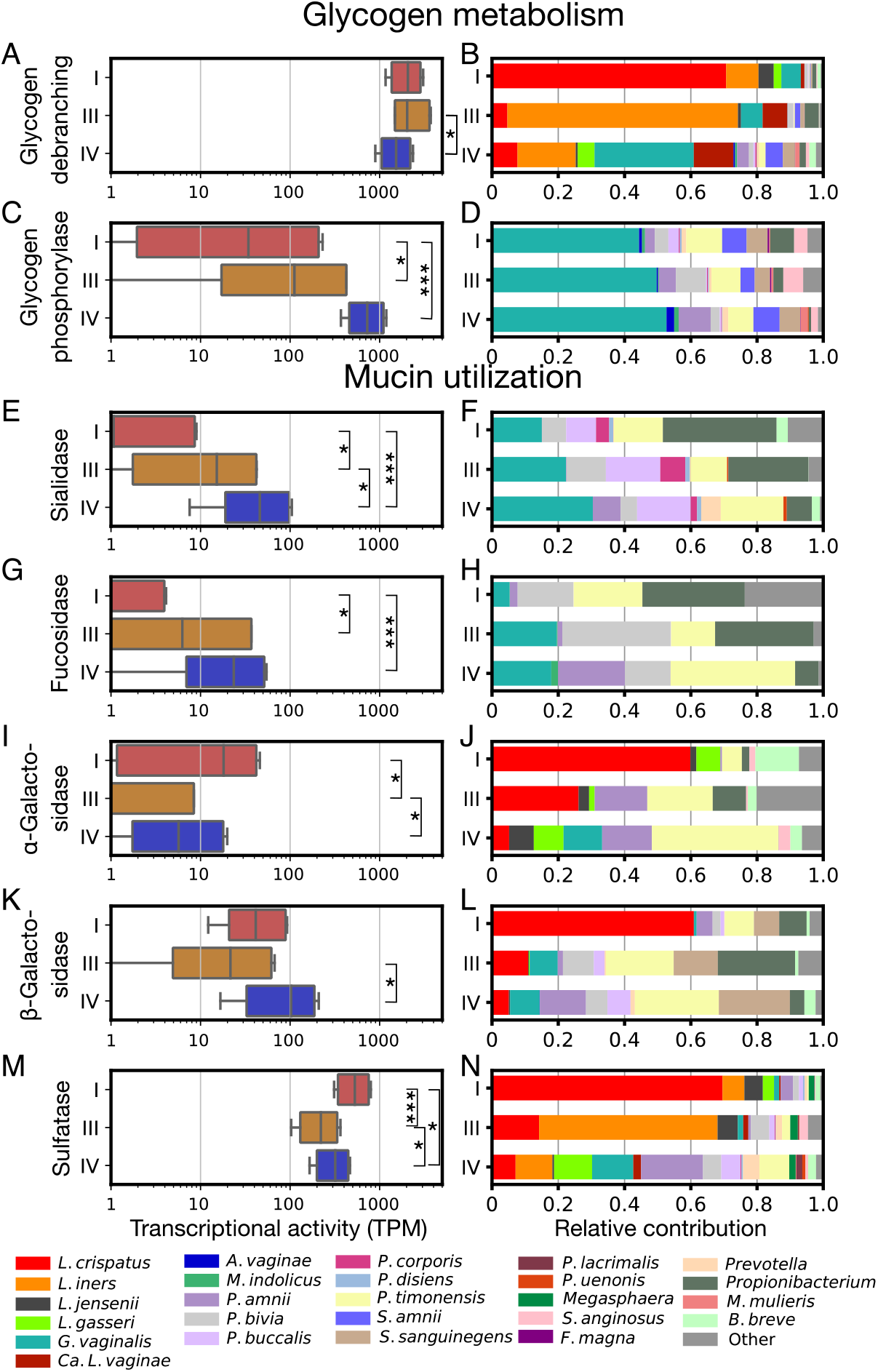
Differences in the expression of key functions related to the metabolism of glycogen and mucin, two abundant host-derived resources in the vagina. Plots on the left (A, C, E, G, I, K and M) display the combined transcriptional output of the specified enzymes in log_10_ transformed transcripts per million (TPM). For each enzyme class, a linear mixed model was used to test for significant differences in its gene expression between CST, with subject included as a random factor. Brackets represent significant *post hoc* comparisons derived from these models. Plots on the right (B, D, F, H, J, L and N) display the average relative contribution of various taxa to the transcription of these genes by members of each CST.

Mucin produced by cervical cells are extensively modified with various carbohydrate residues that, when liberated, can be metabolized by the microbiota (51). The removal of terminal residues, typically sialic acid or fucose, from these mucin-linked oligosaccharides is often the first step in their degradation (52). The enzymes responsible for the removal of sialic acid and fucose are either sialidases (GH family 33) and fucosidases (GH family 29), respectively. The expression of both enzyme families was highest in CST IV communities (Figure 5E and 5F) and lowest in CST I communities indicating that mucin metabolism is likely lower when *Lactobacillus* spp. are dominant in the vaginal microbiota. The taxa responsible for the expression of sialidases in all CSTs include *G. vaginalis*, a number of *Prevotella* spp., along with a minor contribution from *B. breve* (Figure 5F). In comparison, the expression of fucosidase originates primarily from *G. vaginalis, P. amnii, P. bivia, P. timonensis*, and *Propionibacterium* (Figure 5H).

Galactose is another common component of mucin that can be released via the action of either α-galactosidases (GH family 36) or β-galactosidases (GH family 2). Expression of α-galactosidase is higher in CST I and IV communities than in CST III, communities, whereas expression of β-galactosidase is only higher in CST IV as compared to CST III communities (Figure 5I,K). *L. iners* did not express enzymes in either class, leading CST III communities to have the lowest expression of both enzyme families. Non-*Lactobacillus* spp. that express these enzymes include: *G. vaginalis, S. anginosus* (only α-galactosidase), *S. sanguinegens* (only β-galactosidase), and various *Prevotella* spp. Notably, *S. sanguinegens* was responsible for a large proportion of β-galactosidase expression. In sum, these data are consistent with mucin degradation being highest in CST IV communities, with limited degradation in CST I and III because these CSTs lack sialidase and fucosidase expression.

The final class of mucin utilization genes we examined were sulfatases, which act to cleave sulfate groups from mucin, which enables further removal of carbohydrate residues. These enzymes accounted for up to 0.2% of transcriptional activity in communities with their expression being highest in CST I and lowest in CST III (Figure 5M). A wide range of taxa were responsible for their expression including *L. crispatus, L. iners*, and *G. vaginalis* (Figure 5N).

### Integration of metagenomic and metatranscriptomic data identify potential species interactions

In the above analyses we focused on characterizations of the microbiota’s total transcriptional output. Given the high depth of our sequence datasets and the tendency of the vaginal microbiota to exhibit low community evenness, we were also able to examine the transcriptional activity of individual species. The transcriptomes of *L. crispatus* (n=48), *L. iners* (n=74) and *G. vaginalis* (n=58) were found to have adequate coverage (>500,000 reads/genome) in at least 30 metatranscriptomes. We asked whether the gene expression of these species was influenced by the composition of the community in which they resided using an integrative analysis of the metagenomic data to establish community composition and the metatranscriptomic data to define the gene expression of the focal species. Sparse canonical correlation analysis (53) was used to identify axes of covariance between these two datasets, representing combinations of genes from the focal species whose expression were correlated with the relative abundances of combinations of species in the community. To overcome variation in gene sequence and gene content across strains of the same species, we assessed expression at the level of vaginal orthologous protein groups (VOGs) catalogued in VIRGO (34) and removed VOGs which were not present in at least 50% of the samples. The transcriptomic data were also normalized and expressed as transcripts per million reads which corrects for differences in gene length and sequencing depth. The taxonomic composition dataset was filtered to contain only those taxa exceeding a study wide average relative abundance of at least 10^−4^, zeroes were imputed using multiplicative replacement and the data was normalized using the center log ratio (CLR) transformation. In the following paragraphs we describe the results of these analyses and point out aspects of the identified correlations that we find relevant to the ecology of the vaginal microbiota. A complete list of the taxa and VOGs included in each correlation, as well as their weights, can be found in Supplemental Table 3.

For *L. crispatus*, we identified a set of 41 VOGs whose levels of gene expression were correlated with the relative abundance of *A. vaginae* and *G. vaginalis* (ρ=0.80, Supplemental Figure 2A), indicating the abundance of these two species likely influences the transcriptional activity of *L. crispatus*. A principal component analysis (PCA) of the 41 VOGs and two taxa demonstrated that the combined relative abundance of *A. vaginae* and *G. vaginalis* varied along the first axis (Figure 6A). Nineteen of the *L. crispatus* VOGs demonstrated a direct relationship, higher gene expression associated with increasing relative abundances of *A. vaginae* and *G. vaginalis*, and twenty-two VOGs demonstrated an inverse relationship, higher gene expression associated with decreasing relative abundances of *A. vaginae* and *G. vaginalis*. Included among the VOGs with a direct relationship were: two carbohydrate regulatory genes, *glpR* and *cggR*, iron and magnesium transport genes, the *ulaG* vitamin C utilization gene, the alternative sigma factor *sigL*, and the membrane associated *asp23* alkaline shock response gene. VOGs demonstrating an inverse relationship included: the carboxylesterase *nlhH*, the lactose operon repressor *purR*, amino acid and phosphate transporters, the *dnaK* chaperone, and the glutamine synthetase gene *glnA*.

**Figure 6:**
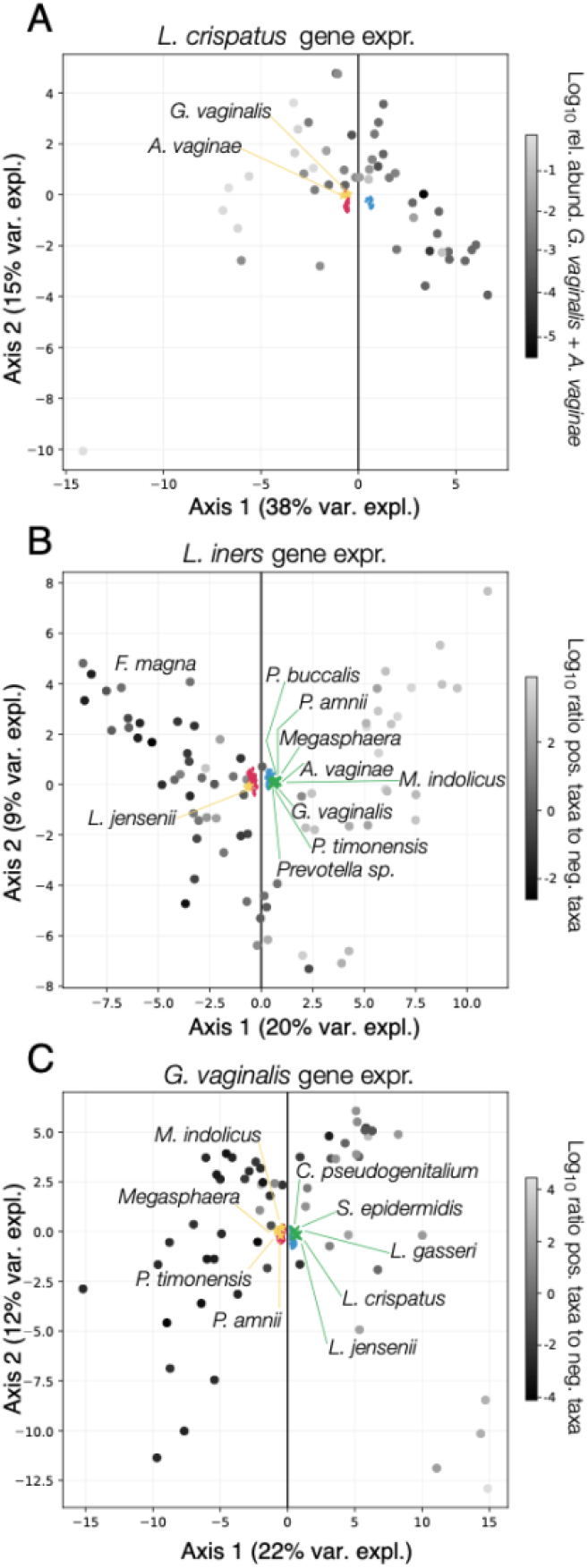
Principal component biplots of the VOGs and taxa identified as correlated by sparse canonical correlation analysis (sCCA). The analysis was conducted on the transcriptional activity of the three most prevalent species: *L. crispatus* (A), *L. iners* (B) and *G. vaginalis* (C). The large circular points in each plot represent the samples. In panel A these points are colored according to the combined log_10_ relative abundance the two negatively contributing taxa (*A. vaginae* and *G. vaginalis*). In panels B and C these points are colored according to the ratio of the combined relative abundances of taxa contributing positively and negatively to the correlation. Factor loadings of the genes (Positive, blue; Negative, magenta) and taxa (Positive, green; Negative, gold) are included in the plot. For legibility, the labels for the taxa loadings, but not the VOG loadings are displayed.

Our analysis of *L. iners* transcriptional activity revealed a set of 111 VOGs which were correlated with the relative abundances of ten taxa (ρ=0.84, Supplemental Figure 2B), two with negative coefficients (*L. jensenii*, and *F. magna*) and eight BV-associated taxa with positive coefficients (*P. amnii, Megasphaera, A. vaginae, P. timonensis, Prevotella spp*., *G. vaginalis, P. buccalis*, and *Mageebacillus indolicus*). This result indicates that the gene expression of these VOGs are correlated with the ratio of positive to negative coefficient taxa and can be seen in a PCA of the combined dataset of correlated VOGs and taxa (Figure 6B). Included among the 58 *L. iners* VOGs associated with a higher relative abundance of *L. jensenii* and *F. magna* was the carbamoyl-phosphate synthase gene (small and large chain components), galactose import and mutarotase genes, the cell shape controlling *rodZ* gene, the cell wall biosynthesis gene *murE*/*murF*, a potassium importer, the NH(3)-dependent NAD(+) synthetase gene, and several components of the DNA damage repair system (*mutL, mutS2, recJ*, and *recR*). On the other hand, the 53 *L. iners* VOGs demonstrating higher expression in the presence of the eight BV-associated organisms included: mannose/fructose/N-acetyl-galactosamine transport genes, the cholesterol-dependent cytolysin inerolysin, a gene encoding a putative mucin-binding protein, glucosamine kinase *gspK*, and the galactose-6-phosphate isomerase *lacAB* genes.

Lastly, we examined the transcriptional activity of *G. vaginalis* using the same approach. Our analysis indicated that a set of 185 *G. vaginalis* VOGs could be correlated with the relative abundances of nine bacterial taxa (ρ=0.80, Supplemental Figure 2C). Of those nine taxa, five were identified as having positive coefficients (*L. crispatus, L. jensenii, L. gasseri, S. epidermidis*, and *Corynebacterium pseudogenitalium*) while four were identified with negative coefficients (*P. amnii, P. timonensis, M. indolicus*, and *Megasphaera*). A PCA of a combined dataset containing the correlated genes and taxa demonstrated that the samples varied in the ratio of the combined relative abundances of the positive and negative taxa along the first axis (Figure 6C). Similar to what we observed with *L. iners*, among the VOGs associated with a higher relative abundance of BV-associated bacteria was the *G. vaginalis* cholesterol-dependent cytolysin vaginolysin. Other VOGs among the 128 associated with a higher relative abundance of these four BV-associated microbes included: a vitamin K epoxide reductase gene, several metal transporters genes, an endopeptidase gene, a serine protease gene, a maltose transport protein gene, and two putative gene encoding glycogen debranching enzymes with alpha-amylase domains. On the other side of the association, we find expression of 57 VOGs to be correlated with higher relative abundances of *L. crispatus, L. jensenii, L. gasseri, S. epidermidis*, and *Corynebacterium pseudogenitalium*. Included among these were *hsp70*, a *groEL* like chaperonin, two genes bearing similarity to the staphylococcal gene *ebh*, two maltose transport genes, two putative glycogen debranching enzymes, and the alkyl-hydroperoxide reductase gene.

## Discussion

Our integrative analysis of vaginal metatranscriptomes and metagenomes has shown that the relative abundance of a species does not necessarily reflect its transcriptional output. Taxa observed at lower relative abundances are often over-represented in the metatranscriptome while taxa observed at higher relative abundances are often under-represented. We have no reason to suspect these differences were due to bioinformatics procedures because the metagenomic and metatranscriptomics datasets were processed in the same manner. We instead posit that this discordance results from differential transcriptional activity among the species in the community. This interpretation has important implications for how we view these communities. If species which we estimate to be rare in the community are highly transcriptionally active, they might have a disproportionate influence on host tissues. For example, *S. amnii* and *S. sanguinigens* were almost always over-represented in the metatranscriptome, and often by several orders of magnitude. These two species are considered by many to be vaginal pathogens and have been shown to be associated with vaginal inflammation *(12)*. Some studies have further linked the abundance of these species to an increased risk of spontaneous preterm birth (36, 54, 55). Associations between these species and adverse health outcomes may be strengthened by examining the transcriptional activity of the community, as the DNA based compositional determinants are likely to underestimate the importance *Sneathia* spp. when present in lower relative abundance.

We found that the transcriptional activity of a species could be used to predict changes in the relative abundance of taxa. Species which were under-represented in the metatranscriptome often decreased in relative abundance by our next timepoint and species which over-represented tended to increase in relative abundance. This result was somewhat surprising given that our timepoints were spaced approximately two weeks apart and that these communities can experience rapid change (31, 32). RNA is less chemically stable than DNA and may therefore more accurately reflect the community at the time of sampling (56). Compositional estimates based on DNA are also likely to experience a lag time while DNA from dead cells is degraded or washed out of the vagina (57). Our results therefore suggest that shifts in community composition are effectuated earlier than our DNA based estimations observe them. This has profound implications for our understanding of the factors that drive changes in the vaginal microbiota. For example, symptomatic bacterial vaginosis may be the manifestation of microbial processes put into motion days prior to its onset. Attempts to discern root causes of BV or to develop novel BV diagnostics would benefit from a closer look at the microbiota’s transcriptional activity before BV manifests.

We next examined functional differences in the transcriptional activity of communities assigned to different CSTs. As defined, communities assigned to different CSTs have fairly distinct taxonomic compositions, so we expected differences in their functional activities. However, functions that differ in expression between the CSTs represent metabolic pathways which could be leveraged to manipulate the composition of the vaginal microbiota, favoring *Lactobacillus*. For example, CST I and III communities express a transporter annotated as being capable of transporting glutamine, glutamate, and aspartate, while CST IV communities express a transporter annotated as only being capable of transporting glutamate. Glutamine is a peptidoglycan precursor and serves as the amine donor for both the conversion of fructose-6-phosphate into glucosamine-6-phosphate and the conversion of aspartate to asparagine in the oligopeptide crosslinks (58). Based on this knowledge, we suggest that glutamine supplementation may favor *Lactobacillus* in the vaginal environment over the BV-associated organisms common to CST IV. This is supported by our observation that *L. crispatus* exhibits increased expression of glutamine synthetase when co-resident with *G. vaginalis* and *A. vaginae*. We also find expression of a metal transporter, predicted to transport zinc and manganese, to be higher in CST I and III than in CST IV. Previous studies have demonstrated that manganese is used by *Lactobacillus* as a defense against oxygen toxigenicity (59), and zinc supplementation has been shown to improve the growth of a probiotic *L. plantarum* strain (60).

Communities dominated by *L. crispatus* were also found to exhibit higher expression of phosphate and phosphonate transporters, consistent with a previous genome comparison that demonstrated *L. crispatus* to have more phosphate/phosphonate transporters than *L. iners* (44) and a previous observation of increased abundance of phosphate transport genes in the vaginal microbiota when vaginal pH is less than 4 (2). Phosphate can buffer pH and intracellular accumulations of polyphosphate have been shown to be used by other *Lactobacillus* spp. as a mechanism of acid tolerance (61). Although *L. crispatus* has not been shown to make polyphosphate, simply sequestering phosphate may allow *L. crispatus* to raise intracellular and lower extracellular pH. Similarly, we found *L. crispatus* dominant communities to exhibit higher expression of a putrescine/spermidine transporter, two biogenic amines capable of buffering pH. This is consistent with the results from metabolomic studies which have found *L. crispatus* dominant communities to have lower levels of putrescine and higher levels of spermidine (27, 62). Manipulating the intracellular and extracellular concentrations of these pH buffering compounds could be a competitive strategy utilized by *L. crispatus* which allows it to inhibit the growth of competitors and facilitate its own metabolism.

As part of our functional analysis, we took a focused look at the expression of enzymes predicted to be involved in the degradation of glycogen and mucin, two host produced potential sources of carbon and energy in the vaginal environment. Glycogen is primarily produced by vaginal epithelial cells (45) whereas mucin is primarily produced by cervical tissues and then secreted into the vagina as a constituent of mucus (46). The ability of the vaginal microbiota to degrade glycogen has been debated for some time (63), but recent studies have demonstrated that many vaginal bacteria likely do have the ability to degrade host produced glycogen (47, 48). We demonstrated that most common members of the vaginal microbiota express predicted glycogen degradation enzymes *in vivo*, although *in vitro* studies are needed to verify their activity. Expression of these enzymes did not appear to vary widely among communities assigned to different CSTs, indicating that glycogen degradation may occur at approximately the same rate in these women and that glycogen is an universal resource in the vaginal microenvironment. Glycogen supplementation alone, which some have suggested to treat BV (64), therefore, seems unlikely to favor the lactobacilli, and might actually be favor the growth of BV-associated bacteria. This challenge might be overcome by complementing glycogen supplementation with a pH lowering agent in order to give the more acid-tolerant *Lactobacillus* an advantage in the metabolism of this common good.

Vaginal mucus has non-specific antibacterial properties and molecules such as lactoferrin, lysozyme, and immunoglobulins are often bound to it (65). Cervicovaginal mucus plays critical functions in fertility (66) and protection against sexually transmitted infections (67), including HIV (68-70). Mucin, the glycoprotein that comprises mucus, appears to be differentially degraded by the vaginal microbiota. Vaginal *Lactobacillus* spp. do not express sialidase or fucosidase enzymes meaning they are likely incapable of removing terminal sialic acid or fucose residues from mucin glycosylation chains. Taxa prevalent in CST IV, including *G. vaginalis* and *Prevotella* spp., do express these enzymes. *Gardnerella* is often featured in discussions of vaginal mucin degradation (49), but our results suggest that *Prevotella* spp., despite typically being at a lower relative abundance in CST IV communities in this cohort, are also likely responsible for a disproportionate amount of this activity. We conclude that the degree and pattern of mucin degradation likely varies with community composition. There have been some direct assessments of variation in vaginal mucin glycosylation chains both through time and between individuals (71). However, these results are difficult to interpret given we know little about variation in mucin composition prior to any microbial tampering. Microbiota driven differences in mucin composition could thus have important reproductive health implications considering the key roles mucins play in fertility and susceptibility to sexually transmitted infections, (68, 72).

In our final analysis we examined the transcriptional activities of *L. crispatus, L. iners*, and *G. vaginalis*. We used an integrative analysis of metagenomic and metatranscriptomic data to ask whether there were sets of genes belonging to each of these species whose expression were correlated with the relative abundances of taxa in the community. The results of these analyses provided functional insight in the ecology of these species in the vaginal environment and hint at possible interactions between members of the microbiota. For *L. crispatus* we find a set of genes whose expression was correlated with the relative abundances of *A. vaginae* and *G. vaginalis*, two species that compete with *L. crispatus* in the vaginal environment. Included in the set of directly correlated *L. crispatus* genes is *asp23*, which encodes an alkaline shock protein. Higher expression of this gene may facilitate the growth of *L. crispatus* under higher pH conditions common to communities with a high relative abundance of *A. vaginae* and *G. vaginalis* (9). Other correlated genes (e.g. carbohydrate metabolism regulators, amino acid and phosphate transporters, glutamine synthase) may signify metabolic shifts in the activities of *L. crispatus* when they co-reside with these bacteria. Additional *in vitro* studies on the shift in transcriptional activity of *L. crispatus* when co-resident with *A. vaginae* and *G. vaginalis* could inform strategies to modulate the microbiota towards an *L. crispatus* dominant state.

The results of our analysis of the *L. iners* and *G. vaginalis* transcriptomes demonstrated a number of commonalities which may reflect similarities in their ecology. *L. iners* and *G. vaginalis* both exhibited differences in their gene expression when co-resident with lactobacilli versus organisms associated with CST IV. For *L. iners*, the set of correlated lactobacilli only included *L. jensenii*, a species which often co-exists with *L. iners* (9). For *G. vaginalis* the correlated lactobacilli included: *L. gasseri, L. jensenii*, and *L. crispatus*. Higher relative abundances of these species were associated with increased expression of alkyl-hydroperoxide reductase by G. *vaginalis*, a protein capable of reducing hydrogen peroxide. *L. crispatus, L. gasseri*, and *L. jensenii* produce H_2_O_2_ *in vitro*, and our transcriptomic data suggests *L. crispatus* expresses enzymes capable of generating H_2_O_2_ *in vivo*. Hydrogen peroxide is toxic to many of the anaerobes common to the vaginal tract, leading many to believe H_2_O_2_ to play a major role in the protection provided by the *Lactobacillus*, although this notion has been challenged (73, 74). Arguments against its relevance include that it is not clear if the generation of H_2_O_2_ is possible at physiological oxygen concentrations in the vagina, and that H_2_O_2_ is unlikely to diffuse far in the vagina far before it is quenched. Our observation is consistent with *Gardnerella* mounting a greater defense against peroxides when co-resident with these *Lactobacillus*, potentially minimizing the impact of *Lactobacillus*-produced H_2_O_2_ and providing another argument against H_2_O_2_ relevance as an antimicrobial. It is also possible that *Gardnerella* is instead responding to elevated environmental O_2_ concentrations which might also favor the lactobacilli. This is an interesting finding, as it suggests that modulating the redox potential of the vaginal microenvironment could be used to favor *Lactobacillus* spp (75).

The second similarity between our analyses of the *L. iners* and *G. vaginalis* transcriptomes using sparse canonical correlation analysis involved their cholesterol dependent cytolysins (inerolysin and vaginolysin (76-78)). Comparative genomic studies of these two species have indicated that both likely acquired their cytolysins via horizontal gene transfer since they are not present in other *Lactobacillus* species or other members of *Bifidobacteriaceae*. In fact, phylogenetic analyses have revealed that inerolysin is most similar to vaginolysin (44), although recent phenotypic comparison have demonstrated some differences between them (79). In both of our analyses, higher expression of the cytolysin was correlated with a greater relative abundance of several BV-associated anaerobes common to CST IV, although only *M. indolicus* was included in both species’ correlations. This result indicates *L. iners* and *G. vaginalis* exhibit higher expression of their cytolysins when in the presence of these organisms, and lower expression when co-resident with other *Lactobacillus* species (*L. jensenii* for *L. iners*; *L. jensenii, L. gasseri*, and *L. crispatus* for *G. vaginalis*). Cholesterol dependent cytolysins act by forming pores in host cell membranes and have been shown to contribute to the pathogenicity of many bacteria (80), although the pathogenicity of the *L. iners* cytolysin is unclear given that the bacterium has not been definitively shown to be associated with adverse health conditions (81). Nevertheless, the pathogenic potential of both *L. iners* and *G. vaginalis* appears to be modulated by the overall composition of the community in which they reside. Communities which are dominated by *L. iners* may not be as destructive to host tissues as those that contain the *L. iners* co-resident with other BV-associated species.

## Conclusion

A common limitation of many microbiome studies is that they rely on DNA-based assessments of community composition. These estimates, be they derived from 16S rRNA gene amplicon sequencing or shotguns metagenomics, represent genomic and functional *potential*. Not all genes on a genome are transcribed at the same time or at the same rate. Sequencing of the metatranscriptome describes the transcriptional activity of microbial communities *in vivo* and give better insight into the function of these communities. However, metatranscriptomics does not provide a complete picture of the biochemical activity of microbial communities. Differences in mRNA stability and translation efficiency can introduce variation in the number of proteins produced per transcript and proteins can vary in their stability. If the transcript encodes an enzyme, its activity will also be governed by the concentration of substrates, products, and cofactors, as well as environmental conditions. Yet metatranscriptomics does bridge a large part of the gap between functional potential and realized biochemical activity. Epidemiological associations between the vaginal microbiome and health outcomes should be strengthened by examining the vaginal metatranscriptome. Our demonstration of the predictive power of the vaginal metatranscriptome could be leveraged for the early diagnosis of vaginal disorders, like symptomatic BV. Further, results from our analysis of metatranscriptomic data could guide the development of innovative strategies to modulate the composition and activity of the vaginal microbiota to restore and maintain an optimal protective environment.

## Acknowledgements

This work was supported by funding from the National Institute of Allergy and Infectious Diseases (UH2AI083264) and National Institute for Nursing Research (R01NR015495) of the National Institutes of Health, and from the Bill and Melinda Gates Foundation (OPP1189217). We also acknowledge the Vaginal Microbiome Research Consortium (VMRC) for helpful discussions.

## Author Contributions

Conceptualization, M. T. F. and J.R.; methodology, M. T. F., B. M., P. G.; formal analysis, M. T. F.; investigation, L. F., L. R., H. Y., M. H., S. N.; writing – original draft, M. T. F.; writing – review & editing, M. T. F., J. R., L. J. F.; visualization, M. T. F.; supervision, J. R., M. H., and L. R.; funding acquisition, J. R and L. J. F.

## Resource availability

### Lead contact

Additional information and requests for additional data and code should be directed to and will be fulfilled by the Lead Contact, Jacques Ravel

### Materials availability

This study did not generate new unique reagents.

### Data and code availability

Metagenomic and metatranscriptomic datasets generated as part of this study have been deposited in the NCBI Short Read Archive under the Bioproject (PRJXXXXXX). Accession numbers for each individual metagenome and metatranscriptome will be found in Supplemental Table 4, upon acceptance and submission. Python and R code used in the described analyses and the generation of figures is available at: (github.com/ravel-lab/XXXX).

## Methods

### Cohort and sample description

The cohort described in this study included 39 North American reproductive-age women, between the ages of 19 and 45. Women included in the study self-identified as Black or African American (n=24), White or Caucasian (n=10), Hispanic or Latino (n=4), and Asian (n=1). They collected vaginal swabs daily, for 10 weeks, according to the protocol described in Ravel, Brotman (41). For each subject we selected up to five timepoints in this timeseries, spaced approximately two weeks apart for metagenomic (n=195) and metatranscriptomic (n=180) sequencing. At time of collection, vaginal swab specimens used in DNA extractions had been resuspended in 1ml of Amies transport medium while RNA extractions had been stored in 2 ml 0.5x RNALater (Thermo Fisher; Waltham, MA) in Amies transport medium—both were preserved at -80°C. Participants also provided behavior and lifestyle information. All studies were performed under Institutional Review Board approved protocols, and samples were collected after obtaining written informed consent from all the participants.

### Shotgun metagenomics

Shotgun metagenomic data was generated for 195 samples (5 samples per subject). DNA was extracted from 200 µL of vaginal swab specimen resuspended in 1ml of Amies transport medium that had been preserved at -80 °C. DNA extractions were performed using the MagAttract PowerMicrobiome DNA/RNA Kit (Qiagen) and bead-beating on a TissueLyser II according to the manufacturer’s instructions and automated onto a Hamilton STAR robotic platform. Shotgun metagenomic sequence libraries were constructed from the DNA extracts using Illumina Nextera XT Flex kits according to manufacturer recommendations and then sequenced on an Illumina HiSeq 4000 platform (150 bp paired-end mode) at the Genomic Resource Center at the University of Maryland School of Medicine. Eight uniquely barcoded samples were included in each HiSeq 4000 lane yielding an average of 35 million read pairs for each sample.

### Shotgun metatranscriptomics

Shotgun metatranscriptomics was generated for 180 samples (maximum of 5 samples per subject, minimum of 2) using a procedure described previously (34). We were unable to obtain quality metatranscriptomic data for fifteen of the samples for which we obtained metagenomics data (information in Supplemental Table 3). For each sample, we extracted bulk RNA extracted from 1.5ml of vaginal swab elute stored in 0.5x RNAlater solution (QIAGEN) that had been stored at -80 °C. Prior to the extraction, 500 µl of ice-cold RNase free phosphate buffered saline (PBS) was added to the 1.5ml of stored swab specimen. To remove the RNAlater, the mixture was centrifuged at 8,000x*g* for 10 min and then the resulting pellet was resuspended in 500 µl ice-cold RNase-free PBS with 10 µl of β-mercaptoethanol. RNA were extracted from the resulting suspension using the Trizol (Sigma) procedure following the manufacturers recommendations. RNA was resuspended in 50 µl of DEPC-treated DNAase/RNAase-free water. Residual DNA was purged from 30µl of RNA extract by treating once with Turbo DNase (Ambion, Cat. No. AM1907) for 30 minutes according to the manufacturer’s protocol resulting in 30µl of RNA suspension. DNA removal was confirmed via a PCR assay using 16S rRNA primer 27 F (5′-AGAGTTTGATCCTGGCTCAG-3′) and 534 R (5′-CATTACCGCGGCTGCTGG-3′). The quality of extracted RNA was verified using the Agilent 2100 Expert Bioanalyzer Nano chip. Ribosomal RNA removal was performed using 9µL of the DNA-free RNA using the RiboZero Plus kit (Illumina) according to the manufacturer’s protocol). Sequencing libraries were then prepared using SMARTer Stranded Total RNA-Seq - Pico Input Mammalian Kit (Takara Bio, Kyoto, Japan) according to manufacturers protocols. The libraries were then sequenced on the NovaSeq platform (150bp, paired-end mode) with 40 libraries included per S2 chip. This provided approximately 85 million read pairs per library. Metatranscriptomic libraries were sequenced deeper than metagenomic libraries to account for the added sequence loss from remaining rRNA.

### Bioinformatic methods

To enable the downstream integration of metagenomic and metatranscriptomic data, the sequence data from both procedures were processed in the same way. First, reads originating from the human host were removed using BMtagger in combination with the GRCh38 human reference genome (82). Ribosomal RNA sequence reads were then removed using sortmeRNA (83). Illumina adapters were excised and reads were trimmed using a 4 bp sliding window with an average quality score threshold of Q15 using Trimmomatic v0.3653 (84). Reads trimmed to a length of less than 75bp were also removed. After these procedures the median number of remaining reads per metagenomic dataset was 2.2*10^7^ (6.6*10^5^-9.9*10^7^) and 1.4*10^7^ per metatranscriptomic dataset (3.0*10^5^-2.0*10^8^). Reads were then mapped to the VIRGO non-redundant gene catalog for the vaginal microbiota using Bowtie (85) using VIRGO default settings (34). Because each gene in the VIRGO database is annotated with a rich set of functional and taxonomic information, the results of these mappings give insight to the function and composition of the vaginal microbiome. To enable our analysis of carbohydrate catabolic enzymes, we further annotated the VIRGO database with the CAZy schema using dbCAN2 (86). These new annotations have been made available via the VIRGO website (virgo.igs.umaryland.edu).

### Analysis of the abundance and activity of taxa

We used the VIRGO mapping results to define the composition and structure of our metagenomic and metatranscriptomic data (Supplemental Table 1). The relative abundance of each taxon was determined by first correcting by dividing the number of mapped reads per gene by its length. These estimates of coverage were combined for each species and divided by sample total coverage in order to compute the relative abundances/contributions. CSTs were assigned based on the taxonomic composition of the metagenome using Valencia (9). For the 180 samples that had both metagenomic and metatranscriptomic data, we computed the relative expression of each taxon by dividing its relative contribution to the metatranscriptome by its relative abundance in the metagenome. We performed two statistical analyses of these taxonomic data as presented in Figure 2. In the first, we modeled the log_10_ relative expression values in response to the log_10_ relative abundance of the taxa at that timepoint (linear mixed model; F_1,4261_=7167, p<0.001), the taxa as a categorical predictor (F_24,4261_=198, p<0.001), and their interaction (linear mixed model; F_24,4261_=11, p<0.001) using a linear mixed model. Subject was also included as categorial random factor to account for our longitudinal data. A second linear mixed model was used to determine whether there was predictive value in our estimates of relative expression. We modeled the log_10_ relative abundance of a taxa as function of the log_10_ relative expression at the previous timepoint (F_1,1847_=649, p<0.001), with taxa (F_24,1847_=11.1, p<0.001) and their combined interaction term (F_24,1847_=3.3, p<0.001) included as well. Tukey adjusted *post hoc* comparisons of the taxa intercepts were conducted for both models.

### Differential expression analysis

Differential expression analyses were conducted between samples assigned to different community state types to identify functional variation in the vaginal microbiome. We conducted these analyses using the KEGG ortholog annotations (42) provided by the VIRGO database because these communities differ dramatically in taxonomic composition. For each KEGG ortholog, we summed the number of reads mapping to genes with the corresponding annotation. We then performed library size normalization using the edgeR package (87) and then modeled the differential expression of each KEGG ortholog using the dream function from the variance partition package in R (88). This package allowed us to fit mixed models which incorporated subject as random factor. Once a differentially expressed KEGG ortholog was identified, we used the VIRGO annotation schema to determine the taxa responsible for its expression. We calculated the relative contribution of each taxa to each differentially expressed KEGG ortholog, and then averaged these values for categories based on which CST they exhibited higher expression in, and the degree of their over-expression. The differential expression analysis was conducted on the metagenomic dataset to determine which KEGG orthologs also showed differential abundance in the microbial community. An additional differential expression analysis was conducted on just the enzymes we predicted using CAZy annotations to be involved in the degradation of glycogen or mucin. For each enzyme, we calculated its normalized expression in transcripts per million, which corrects for differences in gene length and library size. We performed individual linear mixed model analysis with CST as the predictor and subject included as a random factor to test for change in expression. Pairwise Tukey-adjusted post hoc comparisons were made between CST I, III and IV.

### Investigation of species interactions using sparse canonical correlation analysis

We used an integrative analysis of our metagenomic and metatranscriptomics datasets using sparse canonical correlation analysis or sCCA (53) to determine whether the composition of the microbiota influenced the transcriptional activities of *L. crispatus, L. iners*, and *G. vaginalis*. This method identifies axes of covariance between two datasets collected for the same set of samples. We used the individual transcriptomes of these three species as one dataset, and the composition of the microbiota, assessed via the metagenomics data, as the second dataset. First, we identified samples with at least 500,000 reads for the focal species in order to help mitigate differences in library size. Additionally, to help correct for strain differences in the populations of these species, we assessed the transcriptomes at the Vaginal Ortholog Gene (VOG) level (34) representing orthologous genes of the same species. We then removed VOGs that were expressed in less than half of the samples for the three species-specific datasets and converted the read counts per VOG into transcripts per million values. For the compositional dataset derived from the metagenomics data, we took the gene length-corrected relative abundance assessment for these samples. We first removed taxa that demonstrated a study wide average relative abundance of less than 10^−4^ and transformed the relative abundance scores by taking the centered log ratio. To determine whether the pairs of datasets were correlated, we first calculated the RV coefficient of correlation (89) between them using the ade4 R package (90). sCCA was then performed for each of the three species using the PMA package in R (91). Prior to the sCCA analysis, the pairs of datasets were centered and scaled. The permute function was used to select the optimal L1 penalties for both datasets independent. For *L. crispatus*, an L1 penalty of 0.1 was applied to the transcriptomics dataset and a 0.2 penalty was applied to the compositional dataset. These values were 0.2 and 0.4, respectively for *L. iners* and 0.2 and 0.35 for *G. vaginalis*. Final principal components analysis of the correlation VOGs and taxa were conducted using the ade4 package (90).

