## Supplemental Figures for "Insight into the ecology of vaginal bacteria through integrative analyses of metagenomic and metatranscriptomic data"

### Supplemental information

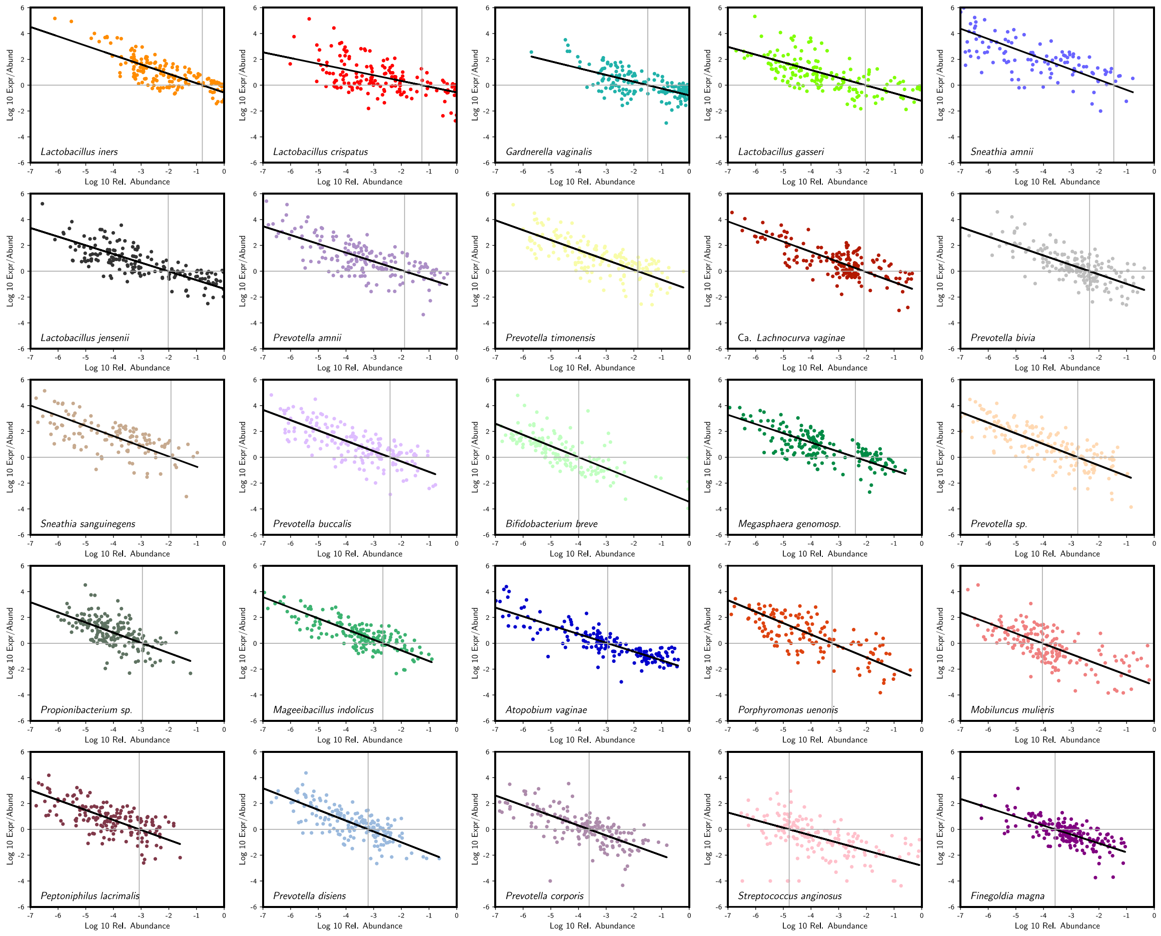


#### Supplemental Figure 1

Linear relationship of the $\log_{10}$ relative expression of each taxa in relation to its $\log_{10}$ relative abundance. Vertical gray lines represent the X intercept of the fit, representing the relative abundance at which each taxon transitions from over- to under-expressive.

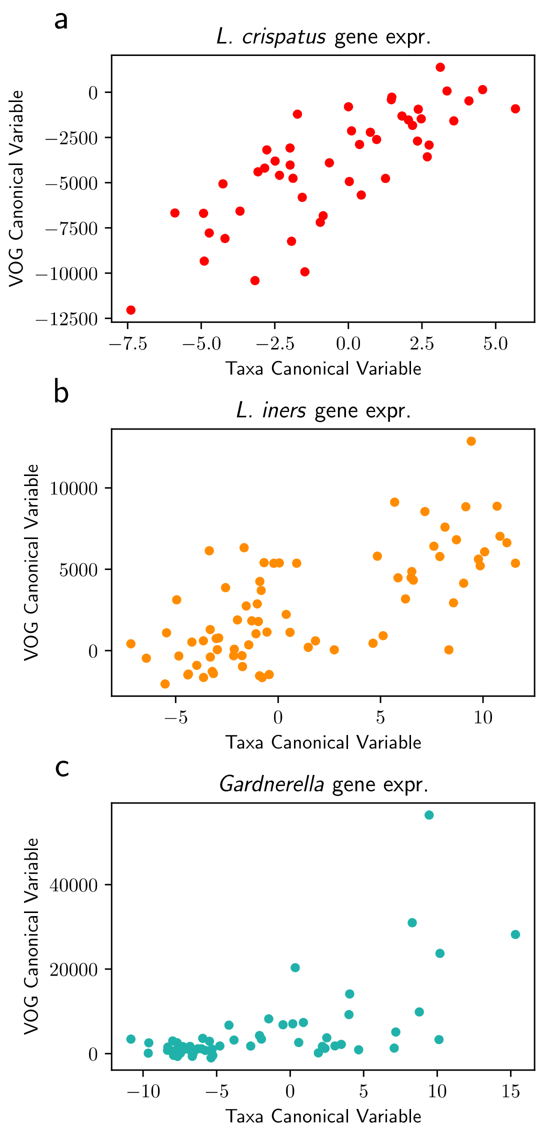


**Supplemental Figure 2**

Scatter plot of correlated canonical variables identified in the analysis of *L. crispatus* (a) *L. iners* (b) and *Gardnerella* (c) gene expression.

**Supplemental Table 1**

Tables containing the taxonomic composition of the metagenomic and metatranscriptomic data, as defined by VIRGO. Taxa relative abundances have been corrected for gene length.

**Supplemental Table 2**

Tables containing results of differential expression analysis at the level of KEGG orthologs

**Supplemental Table 3**

Tables containing non-zero factor loadings for taxa and VOGs identified in sCCA analysis of *L. crispatus*, *L. iners*, *Gardnerella* gene expression.

**Supplemental Item 1**

Graphical displays of the longitudinal taxonomic profile of the metagenomic and metatranscriptomic data for each subject individually.
